# Microscopic and structural observations of actin filament capping and severing by Cytochalasin D

**DOI:** 10.1101/2025.01.28.635382

**Authors:** Takahiro Mitani, Shuichi Takeda, Toshiro Oda, Akihiro Narita, Yuichiro Maéda, Hajime Honda, Ikuko Fujiwara

**Affiliations:** Department of Materials Sciences and Bioengineering, Nagaoka University of Technology, Nagaoka, Niigata, Japan; Department of Complex System Science, Graduate School of Informatics, Nagoya University, Nagoya, Aichi, Japan; Faculty of Health and Welfare, Tokai Gakuin University, Kakamigahara, Gifu, Japan; Department of Biological Science, Graduate School of Science, Nagoya University, Nagoya, Aichi, Japan

## Abstract

Cytochalasin D (CytoD) is widely used to inhibit actin polymerization, but the underlying regulatory mechanism is unclear. We addressed this using Total Internal Reflection Fluorescence (TIRF) microscopy. Our time course depolymerization assay of individual actin filaments showed that CytoD tightly caps the barbed end, with an estimated K_m_ value for inhibition of 4.1 nM and a duration time of ∼1 min. Consistently, in polymerization assays, CytoD at concentrations near the K_m_ value completely suppressed barbed end elongation. Interestingly, at lower concentrations, CytoD acted as a leaky capper, allowing actin monomer addition by rapidly binding to and dissociating from barbed ends. We interpreted this contradictory behavior as arising from differences in binding modes: capping one strand (fast dissociation) or both strands (slow dissociation). CytoD severs actin filaments at micromolar levels, a concentration range commonly used in cell biological studies. Although the severing rate is slower than cofilin, the frequency is higher, resulting in the fragmentation of filaments into shorter pieces. Severing activity was suppressed by inorganic phosphate or cofilin. We determined the crystal structure of CytoD bound to filamentous conformation (F-form) actin and found that CytoD fits better in the hydrophobic cleft of F-form actin than in the monomeric conformation actin, explaining the preferential binding towards barbed end subunits. The structure further indicates that CytoD prevents barbed end depolymerization by stabilizing the terminal subunits in the F-form, which is supported by our MD simulations. Collectively, our results demonstrate how CytoD regulates actin dynamics at the molecular level.

## Introduction

Likely due to the critical role of actin in various cellular processes, including cell motility, cytokinesis, and cell-cell interactions, some primitive organisms produce metabolites as a defense mechanism to target and disrupt the actin cytoskeleton in predators. These actin-targeting compounds, such as phalloidin, latrunculins, and cytochalasins, have been widely utilized in both *in vitro* and *in vivo* actin research. Phalloidin, often conjugated with fluorophores, is a powerful tool for observing the cellular actin cytoskeleton under light microscopy, as it stabilizes actin filaments (1). In contrast, latrunculins and cytochalasins disrupt actin cytoskeleton when added to cell culture media at micromolar concentrations (2–4). Cytochalasins, secondary metabolites of fungi, exhibit a diversity of approximately 400 derivatives (5, 6). Among them, Cytochalasin D (CytoD) is one of the most widely used due to its cell permeability and tight binding to actin (2).

A series of carefully designed biochemical studies, primarily examining bulk actin filaments, have revealed that CytoD acts on actin through the following mechanism. CytoD binds with high affinity (K_d_ =2 nM) to the fast-growing (barbed) end of actin filaments and inhibits both monomer addition and dissociation at that site (7, 8). Although CytoD exhibits only weak binding to monomers (K_d_ = 2 μM) (9), this interaction stimulates ATP hydrolysis activity without inducing polymerization (10, 11). Additionally, CytoD facilitates actin dimerization in a manner dependent on Mg^2+^ and ATP (12, 13).

X-ray crystal structures of monomeric actin with CytoD, obtained through co-crystallization and soaking methods, revealed that CytoD binds to the hydrophobic cleft of actin (11). The occupation of the hydrophobic cleft, exposed only on the barbed end subunits within the filament, likely prevents the interaction with the D-loop of incoming actin monomers, thus explaining the inhibitory effect of CytoD on barbed end elongation. However, the mechanism by which CytoD prevents actin monomer dissociation at this end remains elusive. Furthermore, it is uncertain why CytoD exhibits low binding affinity for actin monomers, despite the hydrophobic cleft being similarly accessible in monomers. The slight flattening of actin observed in the co-crystal structure may provide a clue to the impact of CytoD on actin conformation (11).

The properties of CytoD in barbed end assembly strongly suggest that cytochalasins stabilize intracellular actin networks. However, a report suggested that CytoD treatment leads to the disruption of the actin cytoskeleton (14). In vitro viscosity measurement and electron microscopic observation have indicated the severing activity of cytochalasin B (15, 16). Additionally, elongation assays using acrosomal actin bundles have suggested incomplete inhibition of elongation at the barbed end in the presence of cytochalasin B (17). While these detailed studies have shed light on the destabilizing effects of cytochalasins on actin networks, the molecular mechanisms underlying these effects remain elusive.

Direct observation of individual actin filaments under Total Internal Reflection Fluorescence (TIRF) microscopy revealed that latrunculin A depolymerizes actin filaments by both accelerating monomer dissociation from the terminal end and severing, mechanisms that were previously unidentified (18). This finding prompted us to conduct a comprehensive investigation into CytoD-induced inhibition of actin polymerization. Our TIRF microscopy measurements directly show that CytoD effectively prevents depolymerization at barbed ends for an average duration of ∼1 min. Interestingly, at low concentrations, CytoD functions as a leaky capper, allowing actin monomer incorporation by rapidly binding to and dissociating from the barbed end, consistent with a previous study (17). During the assay, we observed the fragmentation of actin filaments in a CytoD concentration-dependent manner, providing direct evidence of CytoD-induced filament severing. Our crystal structure of CytoD-bound actin in the filamentous conformation (F-form) explains why CytoD binds more stably to F-form actin than to the monomeric conformation (G-form) actin. The conformational preference of CytoD for F-form actin is further supported by MD simulations. Our results suggest that CytoD inhibits barbed end depolymerization by preventing the conformational transition from F-form to G-form at the terminal subunits.

## Results

### CytoD inhibits actin depolymerization from the barbed end

To examine the inhibitory effect of CytoD on actin filament disassembly, we performed a depolymerization assay on individual actin filaments using a TIRF microscope. Actin (1 µM), containing ∼20% AF488-labeled actin, was polymerized in a previously NEM myosin-coated glass chamber. As fiducial marks to measure the lengths to each end of actin filaments, we used the points where the filaments were tethered to the glass surface by NEM-myosin or speckles on the filaments (19). After identifying the barbed end based on its faster elongation rate compared to the pointed end, free actin monomers in the chamber were washed out by loading polymerization buffer with or without CytoD **(as indicated in Fig. 1A and 1B)**. In the absence of actin monomers, typical actin filaments began depolymerization from both ends, with the barbed end depolymerizing much faster than the pointed end **(Fig. 1A)**. In the presence of CytoD, depolymerization from the barbed end was stopped **(Fig. 1B, SI movie 1)**. The plot of filament length confirmed that CytoD specifically blocks actin depolymerization from the barbed end **(Fig. 1C)**. The proportion of capped actin filaments, defined as filaments resistant to depolymerization, increased in a CytoD concentration-dependent manner. At CytoD concentrations exceeding 25 nM, all observed actin filaments stopped depolymerization **(Fig. 1D)**. Note that ∼15% of filaments did not depolymerize even in the absence of CytoD, likely due to nonspecific interactions with NEM-myosin or the glass surface. The estimated K_m_ for CytoD binding to the barbed end was 4.1 nM, consistent with the previously reported value (2). To further investigate the binding duration of CytoD at the barbed end, we measured the time interval between the washout of free CytoD and the onset of depolymerization **(*i.e.*, duration time** τ, **Fig. 1E)**. In standard solution assays, this type of kinetic measurement is challenging due to the difficulty of completely removing free CytoD. To ensure all filaments were capped, we used a sufficiently high concentration of CytoD (25 nM). Higher concentrations of CytoD lead to actin filament severing (see later section). The dissociation rate constant (k^-^_2_) for CytoD from the barbed end, determined from 116 actin filaments, was estimated to be 0.0085 s^⁻¹^ (assuming single CytoD dissociation; see Discussion), corresponding to an average capping duration of 81 seconds **(Fig. 1F)**.

**Figure 1.**
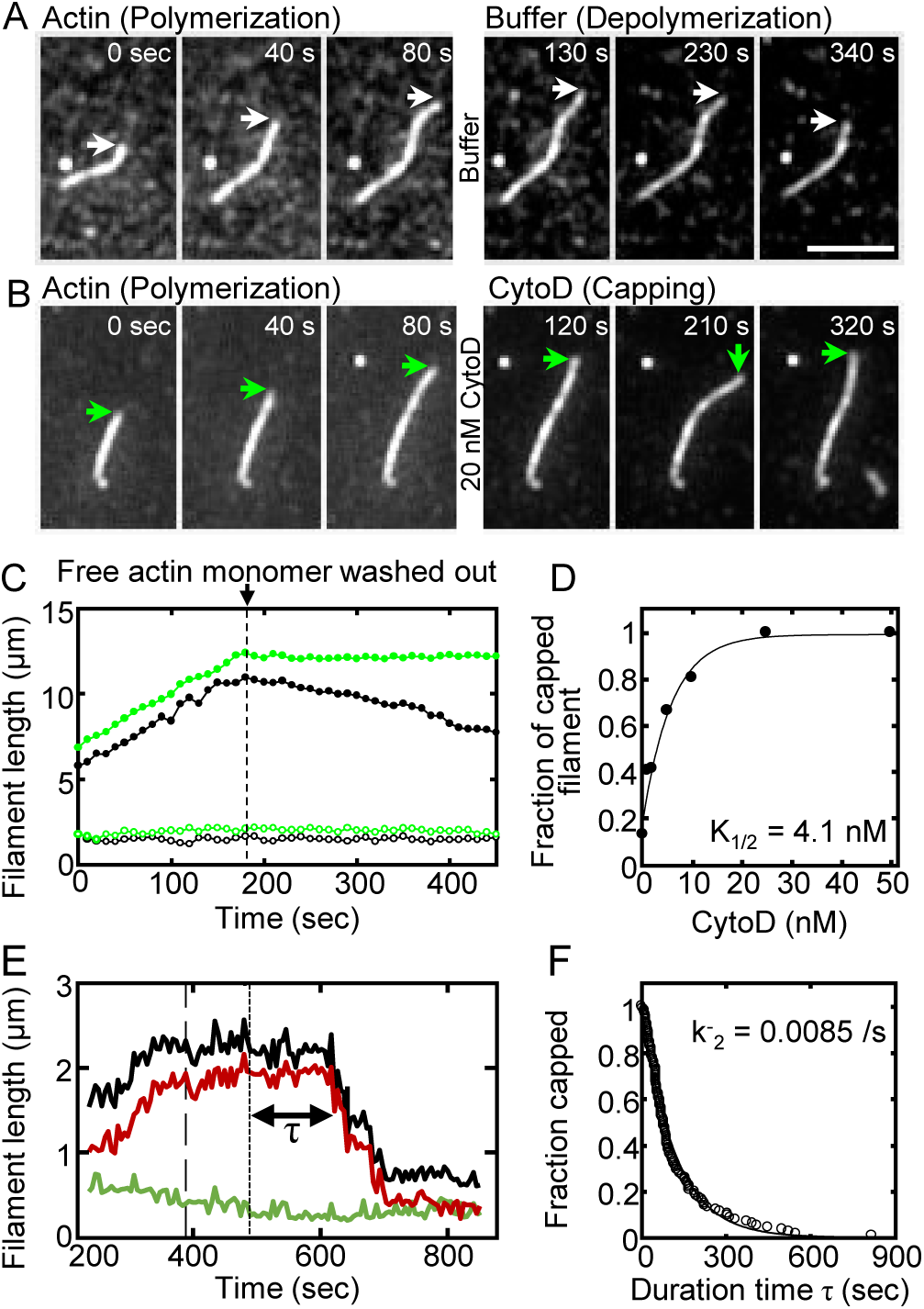
Inhibitory effects of CytoD on actin depolymerization. (A and B) Still images of representative 1 µM actin filaments under TIRF observation over time, containing 16% AF488-labeled actin. The phases of polymerization, depolymerization, or capping are indicated above each phase. Phase changes were initiated by loading the polymerization buffer only (A) or the buffer containing 20 nM CytoD (B) at the indicated time. White (A) and green (B) arrows indicate the barbed ends (fast-growing ends, determined from the polymerization phase) of the filament under each condition. Time zero at the top of each panel marks the start of the recording. Scale bars, 5 µm. (C) Time courses of length changes at the barbed and pointed ends (filled and open symbols, respectively) of typical actin filaments before and after washing out free actin monomer (indicated by the dashed line), in the absence or presence of CytoD (black and green, respectively). (D) Ratio of actin filament that remained capped 500 seconds after the addition of various concentrations of CytoD. The estimated K_1/2_ was 4.1 nM. (E) Length changes at both ends of an actin filament following the addition and subsequent removal of CytoD. The total filament length (black), the length from the fiducial marker to the barbed end (red), and to the pointed end (green) are plotted over time. The dashed line indicates the time of CytoD addition (25 nM), and the dotted line indicates the time of CytoD removal. The duration time is presented as τ. (F) Duration times of capping (τ) for individual actin filaments. The k^-^_2_ value obtained from a single-exponential fit was 8.5 ± 0.2 × 10^-3^ s^-1^, estimated from 116 filaments across two independent experiments.

### CytoD inhibits actin elongation by two modes

To investigate how CytoD inhibits actin polymerization, we monitored the elongation of individual filaments. After confirming the polarity of the filaments from the elongation rate at each end, CytoD, together with actin, was introduced into the chamber. At 5 nM CytoD, which is comparable to the K_m_ value obtained in the previous depolymerization assay, elongation from the barbed end was completely halted **(Fig. 2A and 2C)**. This confirms that CytoD inhibits not only depolymerization but also polymerization at the barbed end, consistent with cell imaging studies showing that CytoD stabilizes short actin filaments at the cell edge through its tight capping function (20, 21). Interestingly, the sustained elongation inhibition was not detected at CytoD concentration below 5 nM; filaments elongated continuously **(Fig. 2B, green)**. The elongation rates were slower than that of free filaments and decreased in a CytoD concentration-dependent manner **(Fig. 2D and 2E)**. Such a slow progressive elongation at barbed ends is reminiscent of the effect found when capping protein (CP) interacts with its allosteric inhibitor, the CAH3 motif of CARMIL (22). CP binds tightly to barbed ends on its own (τ ∼30 min), but its affinity decreases significantly (τ ∼8 s) when bound to CAH3, resulting in rapid association and dissociation of the CP:CAH3 complex at the barbed end. Consequently, in the presence of the CP:CAH3 complex, the elongation profile exhibits a continuous but less steep slope than free filaments due to repeated short pauses that fall below the apparent spatiotemporal resolution of the assay. To determine whether a similar effect occurs with low concentrations of CytoD, we plotted the length change of individual actin filaments from their barbed ends every 60 seconds (in bin sizes of approximately 160 nm, the same resolution of the microscope used in this assay) across different CytoD concentrations **(Fig. 2F)**. If the slowing of elongation is driven by the rapid association and dissociation of CytoD, the histograms should represent the sum of two phases originated from elongation and pause. Global fitting confirmed the presence of two Gaussian distributions: one (red distribution) with a median rate of 7.1 sub·s⁻¹ (1.1 μm·min⁻¹), which matches the expected elongation rate for 0.7 µM actin, and another (blue distribution) with a median rate of 1.3 sub·s⁻¹ (0.2 μm·min ¹), corresponding to the pause phase. The rate representing the pause phase is not zero, likely due to the relatively coarse spatial resolution of our measurement, 0.16 µm·pixel⁻¹. The ratio of the pause phase increased as the CytoD concentration increased. Therefore, we conclude that at low concentrations, CytoD binds to the barbed end only transiently. In summary, our single filament observations revealed two modes of CytoD-induced elongation inhibition at barbed ends: at high concentrations, CytoD acts as a tight capper, whereas at low concentrations, it behaves as a leaky capper, rapidly associating with and dissociating from barbed ends, thereby allowing slow monomer association.

**Figure 2.**
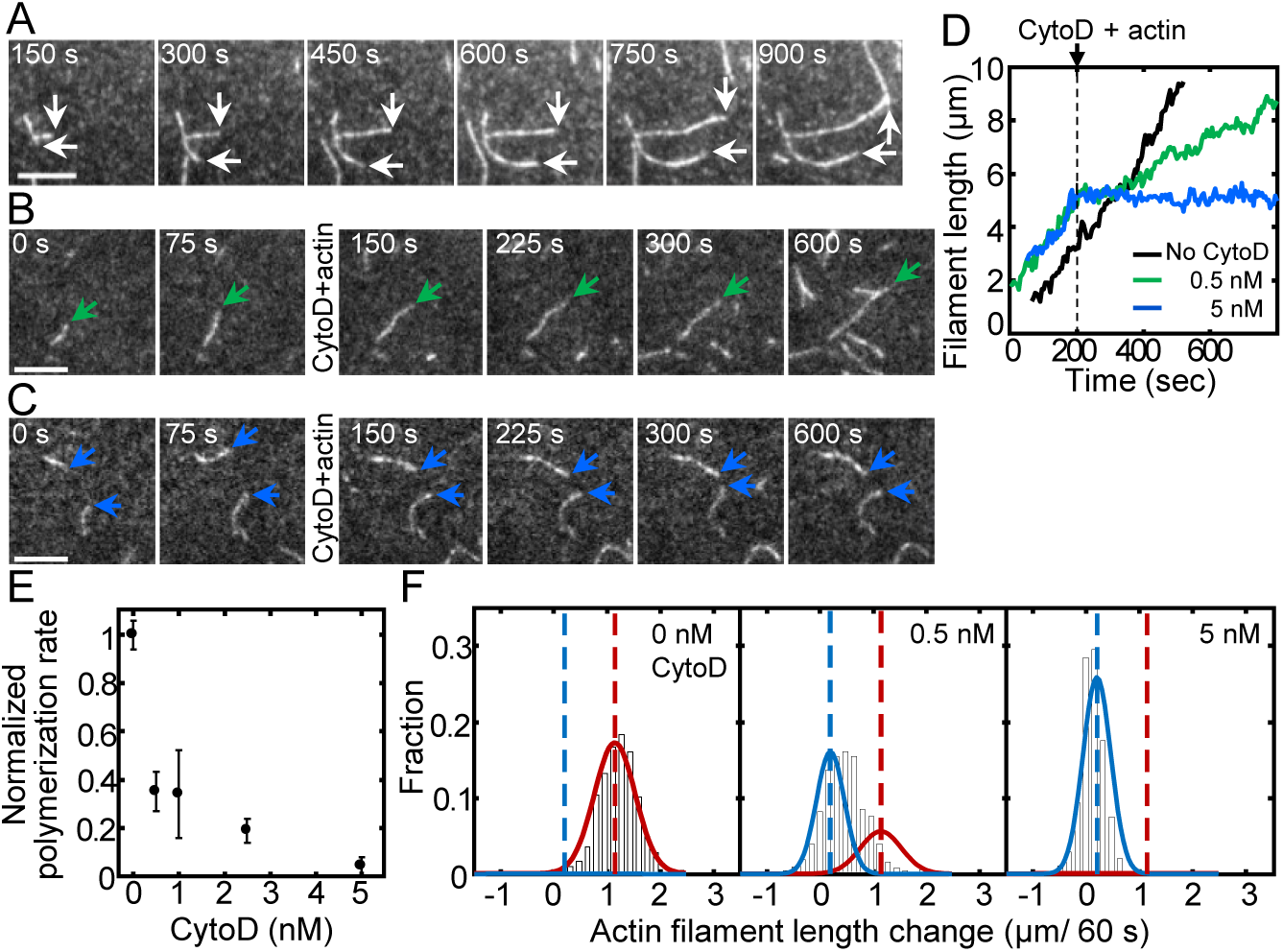
Inhibitory effects of CytoD on actin polymerization. (A-C) Time-lapse TIRF images of actin polymerization (0.7 μM with 20% AF488-labeled) in the absence (A) and presence of 0.5 nM (B) and 5 nM (C) CytoD. Arrows indicate the barbed ends of actin filaments. CytoD, together with 0.7 µM actin in polymerization buffer, was loaded into the chamber 75 seconds after initiating polymerization, as indicated time in the image. Scale bars, 5 µm. (D) Time-course plot showing the length of representative actin filaments in the presence of 0 (black), 0.5 (green), and 5 nM (blue) CytoD, with 0.7 µM actin in the polymerization buffer. The dashed line indicates the time of CytoD addition. (E) Decrease in the apparent elongation rate of the barbed end as a function of CytoD concentration. Each data point represents the average of 10 actin filaments from three independent experiments. (F) Histograms showing changes in the barbed end length of ∼10 actin filaments every 60 seconds. The concentrations of CytoD (0, 0.5, and 5 nM) are indicated in the top right corner of each panel. Two Gaussian distributions, globally fitted, show peaks at 1.1 and 0.2 µm/min (corresponding to 7.1 (red) and 1.3 (blue) subunits/sec, respectively).

### CytoD Severs Actin Filaments

Interestingly, we often observed the breakage of actin filaments in the presence of high concentrations of CytoD **(Fig. 3A)**. Although previous studies (2, 3) have suggested a severing function for CytoD, their investigations were limited by technical difficulties. Using our current approach with TIRF microscopy, we were able to directly observe the severing of individual filaments. To eliminate actin monomer association, which potentially complicates the analysis, we observed the severing reaction under depolymerization conditions. An order of magnitude higher concentration of CytoD was required to induce severing of actin filaments compared to capping the barbed end, indicating that CytoD has a lower affinity for binding to the sides of actin filaments than to the terminal subunits. Under these conditions, actin depolymerization from the barbed end was also suppressed, leading to a stepwise decrease in filament length **(Fig. 3B)**.

**Figure 3.**
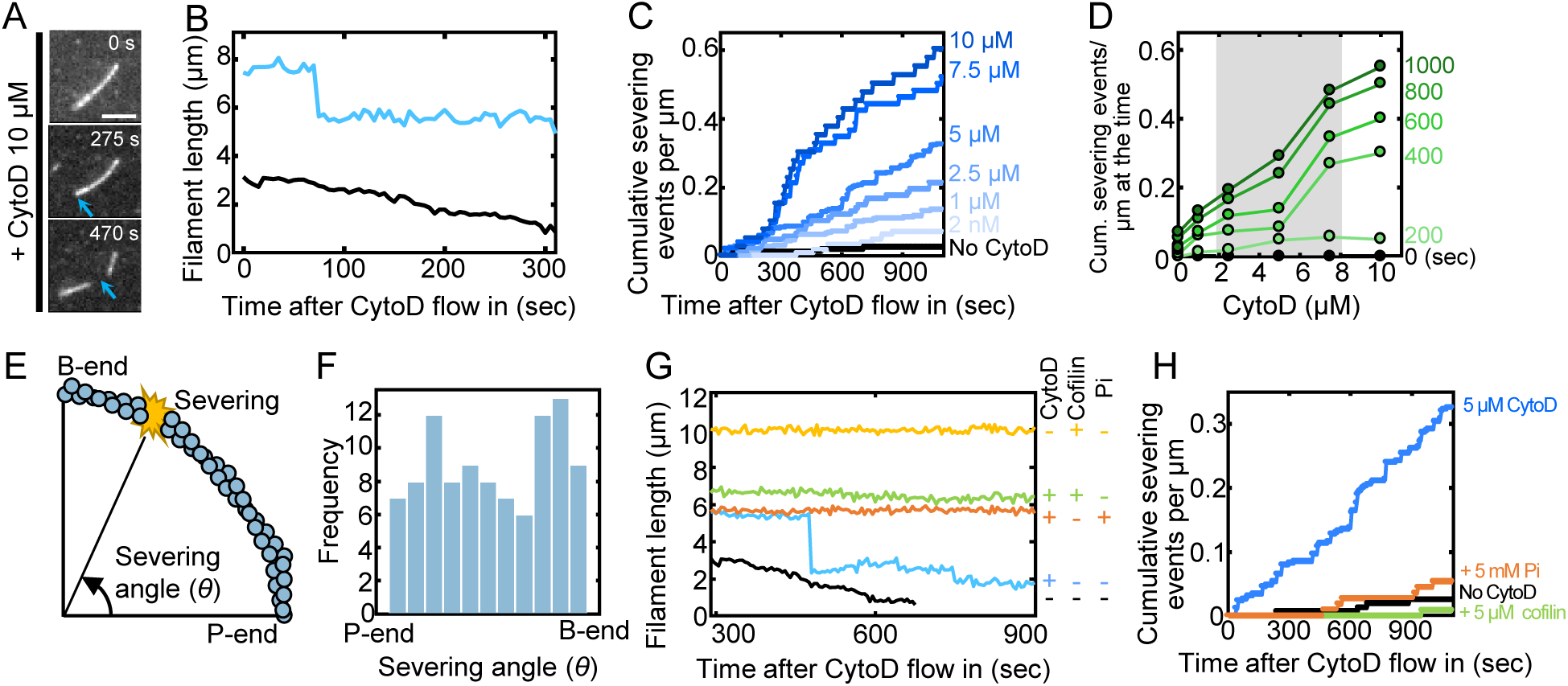
CytoD severs actin filaments. (A) Time-lapse images showing actin filament severing following the addition of CytoD. The time after CytoD addition is indicated in the top right corner of each panel. Blue arrows mark severing events. Scale bar: 5 µm. (B) Length change of an actin filament severed by 10 µM CytoD (blue). The length change of an actin filament without CytoD is shown in black for comparison. (C) Cumulative severing events on actin filaments per µm over time after loading various concentrations of CytoD. The concentrations of CytoD are indicated on the right side of the panel, with colors corresponding to the data. (D) Cumulative severing events within 0 to 1000 seconds after loading various concentrations of CytoD. Data were collected from 20-80 actin filaments across 2-3 independent experiments. The gray area indicates CytoD concentrations typically used in cell biological research. (E) Schematic illustration showing the method used to determine the location of severing along actin filaments. (F) Frequency distribution of CytoD-induced severing location along actin filaments. (G) Temporal length change of actin filaments in the absence (black) or presence of 5 µM CytoD (blue) with 5 mM P_i_ or 3 µM cofilin (orange and green, respectively). The length change of an actin filament in the presence of cofilin without CytoD is also plotted (yellow). (H) Cumulative severing events on actin filaments in the absence (black) or presence of 5 µM CytoD (blue) with 5 mM P_i_ or 3 µM cofilin (orange and green, respectively).

To investigate the severing kinetics, we counted cumulative severing events, defined as severing frequency per unit length (1 μm) of actin filaments **(Fig. 3C)**. Over a 1200 second observation time frame, severing events became more prominent at 5 μM CytoD and reached saturation at 7.5 μM CytoD. This concentration dependence confirmed that severing was indeed caused by CytoD, (not by mechanical shear force in the solution). Our results showed that approximately six severing events occurred in a 10 μm long actin filament (see 0.6 on the y-axis). A similar experiment indicated that cofilin-induced severing saturated at only 0.2-0.3 events per µm (23). Cofilin severs filaments through cooperative cluster binding over a certain length, whereas such binding is unlikely to be involved in CytoD-mediated severing, which may explain CytoD’s higher efficiency. Notably, the time course of severing events followed a sigmoidal curve, with an approximately 200 second lag phase before the initial severing, even at high concentrations **(Fig. 3C and 3D)**. This also contrasts with the cofilin-induced severing experiment, which exhibited a shorter lag time (23). A possible explanation is that CytoD would preferentially sever ADP-bound actin filaments (*i.e.*, aged filaments), and under our experimental conditions, the filaments were not well-aged when CytoD was added. Next, we mapped the spatial distribution of severing points along actin filaments **(Fig. 3E and 3F)**. The plot showed that severing takes place more readily near the ends of filaments than in the middle, which may be due to the larger structural fluctuations inherent to the filament ends.

### Cofilin and P_i_ Suppress CytoD-induced Severing

CytoD has been reported to inhibit cofilin binding to actin filaments (14), which, in turn, suggests that cofilin-decorated actin filaments (cofilactin filaments) hinder CytoD-induced severing. To investigate this hypothesis, we quantified the severing events of cofilactin filaments in the presence of CytoD **(Fig. 3G and 3H, Fig S1A and S1B)**. To prepare cofilactin filaments, bare actin filaments were incubated with cofilin in the flow chamber at a slightly acidic pH of 6.6 to avoid cofilin-induced filament fragmentation (24). Under the acidic condition, CytoD still severed bare actin filaments **(Fig. S1A)**, similar to the assays conducted at pH 8.0 **(Fig. S1C)**. In contrast, cofilactin filaments remained mostly intact **(Fig. S1B and S1E)**, demonstrating that CytoD-induced severing is blocked by cofilin.

We next examined the severing activity of CytoD on individual actin filaments in the presence of inorganic phosphate (P_i_) **(Fig. 3G and 3H, Fig S1C and S1D)**. Actin subunits hydrolyze bound ATP to stabilize the filamentous structure, with ADP-P_i_-bound actin being the most stable state (25, 26). Therefore, we investigated whether CytoD could sever ADP-P_i_-actin filaments, which were prepared by adding 5 mM P_i_ to the solution. We found that P_i_ suppressed CytoD-driven severing **(Fig. S1C and S1D)**; the cumulative severing events by CytoD on ADP-P_i_-actin filaments were almost indistinguishable from the control (no CytoD) **(Fig. 3H)**. Our results indicate that, like cofilin and gelsolin, CytoD is ineffective at severing ADP-P_i_-actin filaments.

### Crystal structure of CytoD bound to F-from actin

CytoD preferentially binds to actin subunits at the barbed end adopting the F-form (K_d_ in the nanomolar range) (7, 8 and this study) rather than to monomeric actin adopting the G-form (K_d_ in the micromolar range) (9), even though the binding site for CytoD, the hydrophobic cleft, is exposed in both cases. To address the structural basis for this conformational preference of CytoD, we aimed to obtain the missing information, *i.e.*, the CytoD-bound F-form actin structure. We utilized the F-form actin crystallization system (F1A) in which the N-terminus domain of fragmin (F1) binds to an actin molecule (A) and fixes it in the F-form (27)**(Fig. S2C)**. The flat conformation of actin is maintained by the N-terminal extension of fragmin (FNEX), where Asn13 forms extensive interaction with actin’s Pro-rich loop (residues 108-112). Our attempt to soak the F1A crystal with CytoD failed, likely due to competition with FNEX, which also targets the hydrophobic cleft (**Fig. 4E and** see below). To facilitate CytoD binding by reducing the affinity of FNEX for actin, we replaced Asn13 with Ala. Although the incubation of ATP-actin with F1^N13A^ did not yield crystals, the addition of CytoD to the proteins produced a crystal that diffracted to 1.7 Å resolution. The refined structure, F1^N13A^A_CytoD, consists of F1^N13A^, actin, and CytoD with full occupancy positioned between the inner domain (ID; subdomains 3 and 4) and outer domain (OD; subdomains 1 and 2) **(Fig. 4A-C and Fig. S2A)**, as observed in two CytoD-bound G-form actin structures determined by soaking (3EKU) or co-crystallization (3EKS) (11). As anticipated, the binding site for FNEX partially overlaps with CytoD at the back half of the hydrophobic cleft, resulting in the complete dissociation of FNEX **(Fig. 4E)**, a prerequisite for maintaining the F-form actin conformation in the F1A crystals **(Fig. S2C and S2D)** (27). Yet, the actin conformation in F1^N13A^A_CytoD remains F-form, with the bound ATP completely hydrolyzed to ADP-P_i_ **(Fig. 4D and Fig. S2E)**, indicating that, instead of FNEX, CytoD is responsible for stabilizing the F-form actin conformation in the present crystal. The capability of CytoD to induce actin flattening was suggested by Nair and colleagues **(Fig. S2F-H)** (11). Compared to their CytoD-soaked structure, actin exhibited a slightly flatter conformation (2.2°) in the co-crystallized structure **(Fig. S2G and S2H)**, where the protein conformation is assumed to be free from the constraints of the original crystals.

**Figure 4.**
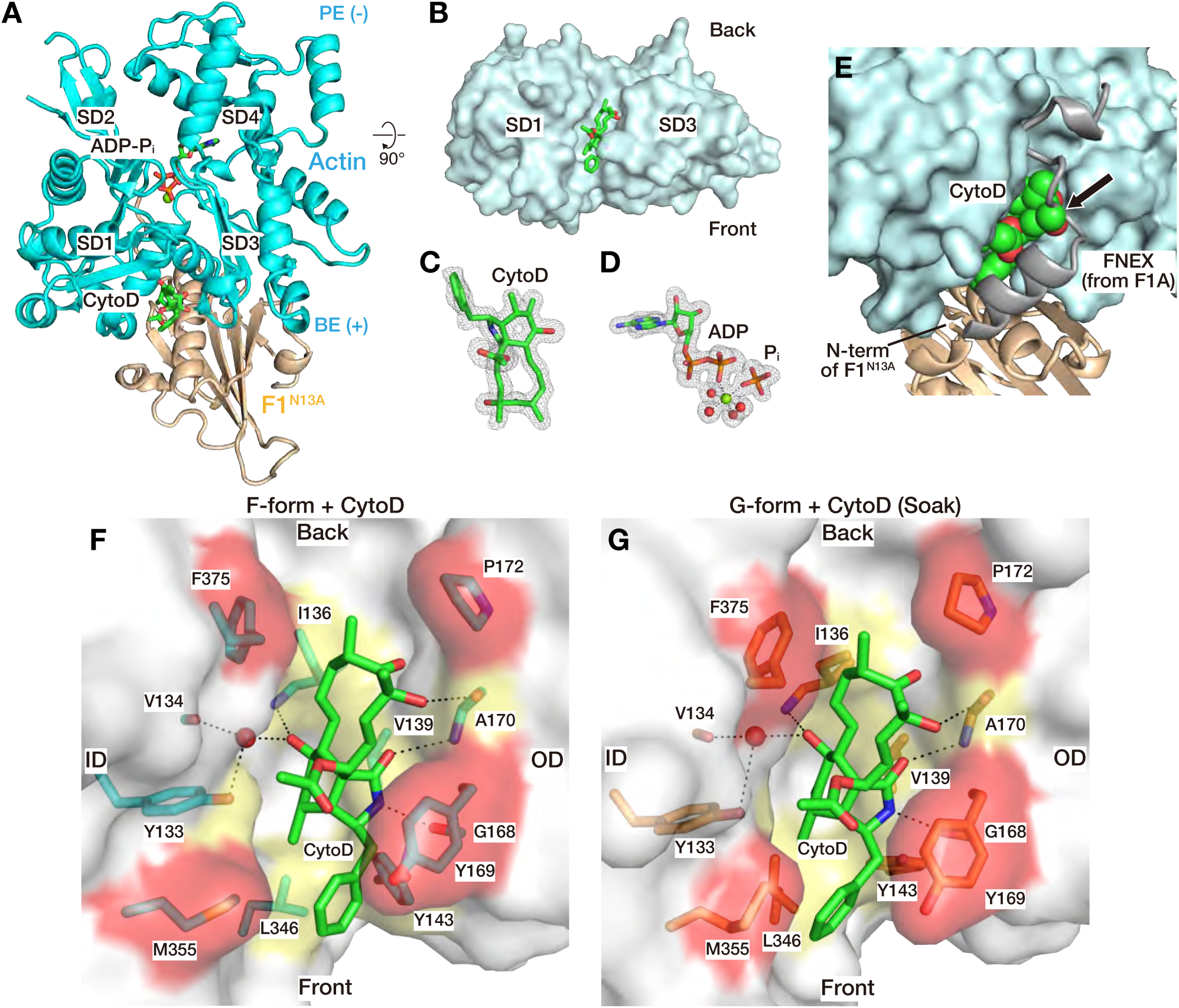
Crystal structure of CytoD-bound F-form actin. (A) Overall structure of F1^N13A^A_CytoD. This and the following panels show actin viewed from the back side, opposite to the standard presentation by (64). Actin subdomains (SD1–SD4) are labeled for reference. (B) Bottom view highlighting the position of CytoD in the hydrophobic cleft. F1^N13A^ is omitted for clarity. (C and D) *Fo - Fc* omit maps (contoured at 4 σ) for CytoD (C) and ADP-P_i_ (D). Green and red spheres represent Mg^2+^ and its coordinated water molecules, respectively. (E) Overlap between CytoD and FNEX. FNEX does not bind to actin in F1^N13A^A_CytoD. The N-terminus residue of F1^N13A^ present in the structure is Glu28. FNEX (gray), derived from the F1A structure without CytoD (PDB 7W50), is modeled onto F1^N13A^A_CytoD. A black arrow indicates a clash between CytoD and the backbone of FNEX residues Asn16-Gly18. (F and G) CytoD binding site in F-form actin (F; this study) and G-form actin (G; soaking crystal 3EKU). CytoD and its binding actin residues are depicted as sticks, with hydrogen bonds shown as dotted lines. Water molecules mediating CytoD binding are depicted as red spheres. Actin is shown in white surface representation, with CytoD-contacting residues colored yellow (buried surface area upon binding to CytoD >15 Å^2^) and red (>30 Å^2^).

Despite CytoD’s preference for F-form actin, the mechanism of ligand recognition is highly similar between G-form and F-form actin structures **(Fig. 4F and 4G)**. In both structures, the same set of actin residues forms hydrogen bonds with CytoD, some mediated by water molecules. Similarly, CytoD, deeply embedded in the hydrophobic cleft, forms van der Waals contacts with actin residues common to both the G-form and F-form **(Table S2)**. Notably, residues Tyr169, Pro172 (ID), Met355, and Phe375 (OD) at the bottom of actin contribute significantly to CytoD binding, as evidenced by their substantial buried surface areas upon CytoD binding (>30 Å²). Taken together, our inspection of the ligand binding site suggests that, while minor differences might have an impact, the flattening of actin does not essentially create additional contacts that would dramatically increase its binding affinity for CytoD.

Why, then, can CytoD bind stably to F-form actin? A comparison of F1^N13A^A_CytoD with F1A shows that the CytoD binding does not induce major structural change in actin (Cα RMSDs = 0.13 Å), except for the C-terminus residue Phe375, which partially occupies the hydrophobic cleft in the CytoD-free structure **(Fig. S3A)**. Upon CytoD binding, the entire Phe375 residue, including its main chain, flips upward towards the Pro-rich loop **(Fig. S3B)**. The repositioned Phe375 is stabilized by polar interactions with Arg116 and His371, as well as by nonpolar interactions with Ile136 and the macrocycle of CytoD (C13-C16) **(Fig. 5A and Fig. S3B;** the chemical structure of CytoD is shown in **Fig. S2B)**. Thus, the flipping of Phe375 is not only to avoid CytoD but is essential for creating a hydrophobic pocket that accommodates CytoD **(Fig. S3E)**. No major steric clashes are observed in this site **(Fig. 5D)**. Phe375 also undergoes a CytoD-induced shift in G-form actin structures **(Fig. S3C and S3D)**; however, this causes numerous steric clashes **(Fig. 5C and 5F)**. For example, in the soaking structure, Phe375 is only 2.7 Å away from Phe375, which is too short for atoms that do not form polar contacts. This is due to the downward shift of the Pro-rich loop in G-form actin compared to F-form actin. The substantial structural instability around Phe375 accounts for the low affinity of G-form actin for CytoD. In the co-crystallization structure (3EKS), although Phe375 remains close to Glu107 (2.8 Å), the clashes around Phe375 are largely alleviated **(Fig. 5B and 5E)**, supporting the idea that the flattening conformational change of actin is favorable for CytoD binding. It is likely that CytoD readily binds to the actin subunits at the filament barbed end, as observed in our F-form actin structure **(Fig. S3F-S3J)**. In conclusion, optimal CytoD binding is achieved at the hydrophobic cleft of F-form actin. In G-form actin, the slightly different ID/OD arrangement compared to F-form actin causes local distortions around Phe375 upon CytoD binding, leading to weaker affinity.

**Figure 5.**
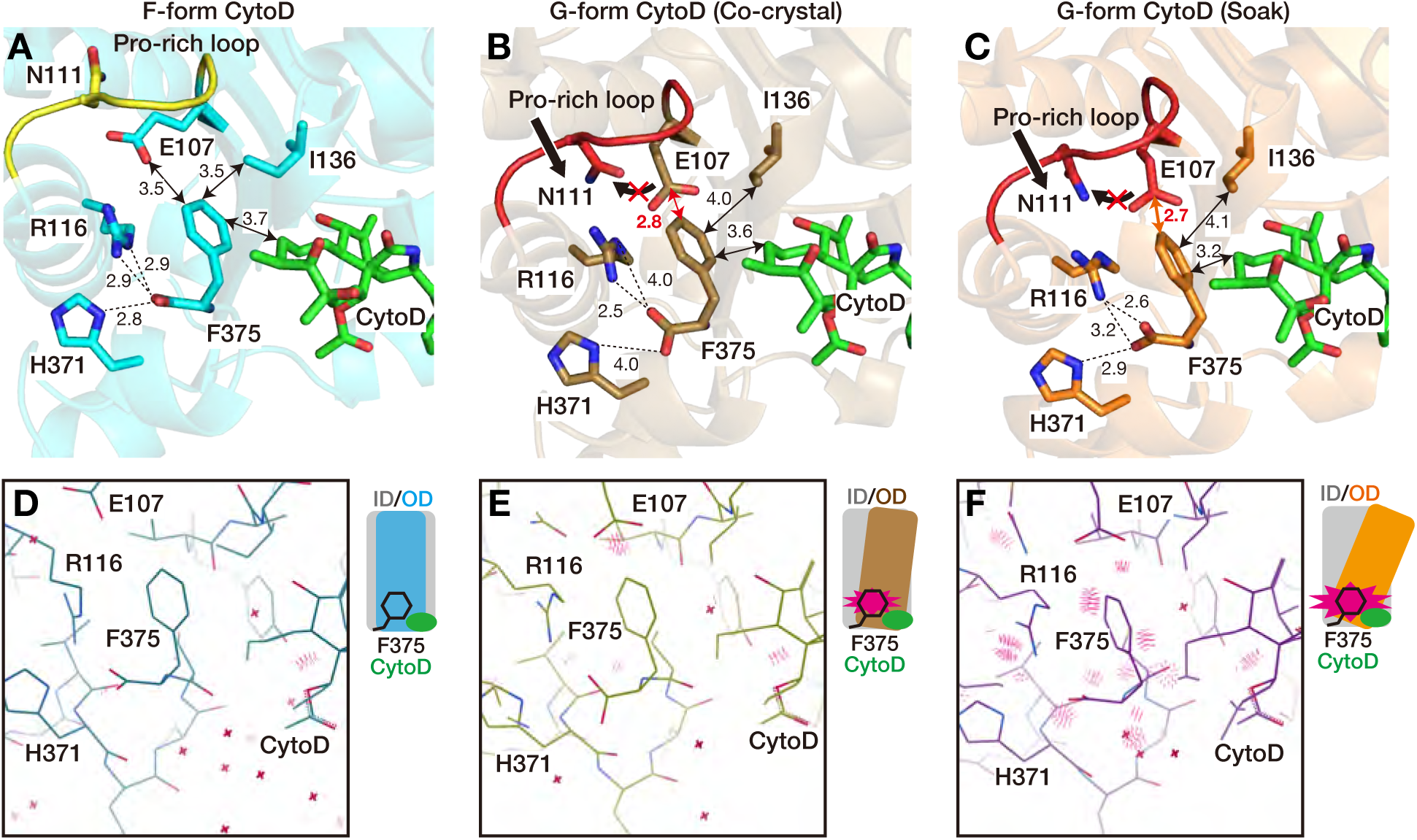
Positioning of Phe375 in CytoD-bound actin. (A-C) The environment surrounding Phe375 in F1^N13A^A_CytoD (A), CytoD-bound G-form actin (co-crystallized; 3EKS) (B), and CytoD-bound G-form actin (soaked; 3EKU) (C). Key residues interacting with Phe375 are shown as sticks. Hydrophilic and hydrophobic interactions with normal (non-clashing) distances (Å) are indicated by black dotted and black double-arrowed lines, respectively. Red double-arrowed lines represent sterically unfavorable close contacts between Phe375 and Glu107. In F1^N13A^A_CytoD, the Pro-rich loop (residues 108-112; yellow) is positioned away from the hydrophobic cleft, and the Glu107 side chain extends upward, maintaining a 3.5 Å distance from Phe375, with Asn111 oriented oppositely. In G-form actin, the Pro-rich loop (red) shifts downward compared to the F-form structure, causing a sterically unfavorable proximity of the Glu107 side chain to Phe375 (2.7-2.8 Å). Asn111 in the Pro-rich loop of G-form actin limits the orientation of Glu107, preventing it from avoiding Phe375. (D-F) Steric clashes around Phe375 in CytoD-bound F-form actin (D), CytoD-bound G-form actin (co-crystallized) (E), CytoD-bound G-form actin (soaked) (F). Steric clashes were evaluated using Probe (65) implemented in Coot (55). Severe clashes (steric overlaps larger than 0.9 A) are indicated by pink spikes, with their size representing the degree of overlap. Notably, clashes observed in the soaked structure are largely resolved in the co-crystallized structure, suggesting that actin flattening is favorable for CytoD binding.

### MD simulation of CytoD-bound actin

To further investigate CytoD’s preference for actin conformation, we performed MD simulations. After removing the fragmin F1 from F1^N13A^A_CytoD and optimizing the F-form actin structure (ADP-P_i_ state) with and without CytoD, three 400 ns runs were conducted for both structures. The time course of OD orientation relative to ID of actin was measured, using the ADP-G-actin structure (1J6Z (28)) as a reference for the standard orientation (0°). The G/F axis angles, representing the degree of flattening between the two domains, is typically 14-16° in F-form actins **(Fig. 6A)** (29). In the absence of CytoD, the G/F angle decreased from ∼14° towards 0° in all three runs **(Fig. 6B)**, consistent with the instability of the F-form conformation, even in the presence of P_i_ in the active site. In contrast, with CytoD present, the G/F angle remained relatively unchanged throughout the simulations **(Fig. 6C)**, indicating that CytoD binding stabilizes actin in the F-form. Both CytoD (this study) and F1 with intact FNEX (F1^1-160^) (30) stabilized actin in the F-form in MD simulations. However, CytoD-bound actin exhibited larger deviations from the initial structure (black points in the graph) during the simulations, with more pronounced changes along the G/C axis, which is perpendicular to the G/F axis **(Fig. 6D and 6E)**. CytoD-bound actin tends to feature a more open nucleotide-binding cleft **(Fig. 6F)**, which might have a potential impact on inter-actin subunit interactions.

**Figure 6.**
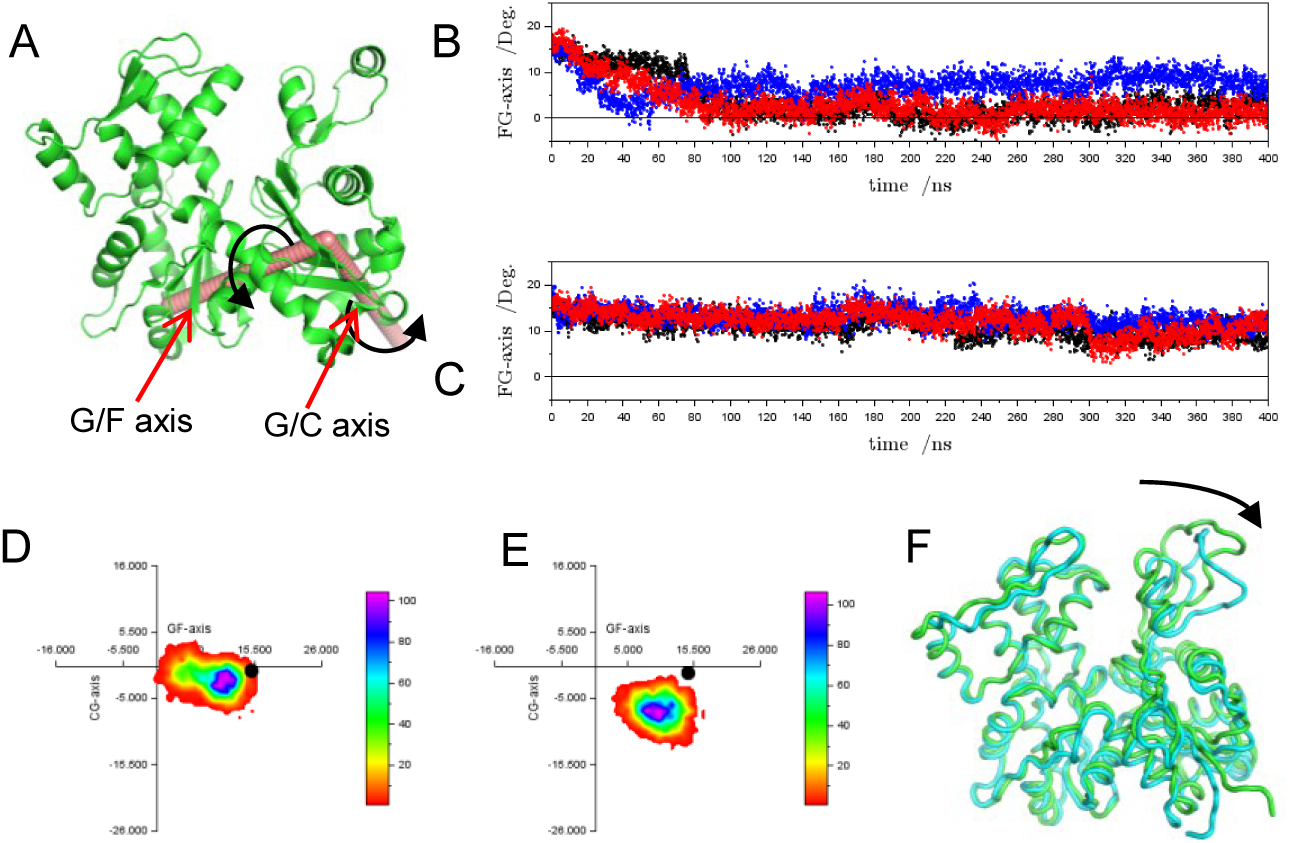
MD simulation of CytoD bound to F-form actin. (A) G/F axis and G/C axis shown in actin (PDB code: 1J6Z). The rotation angles, GF-axis angle and GC-axis angle, around the axes were used to analyze of the orientation of OD relative to ID. The arrows indicate the directions of rotation. The GF-axis angle and GC-axis angle represent the degree of flattening of the two domains and the closure of the nucleotide-binding cleft, respectively. (B) Time course of GF-axis angles during the MD simulations of actin alone. The initial structure was a crystal structure of F1^N13A^A_CytoD, with F1^N13A^ and CytoD removed. The colored scatter points represent different independent runs. (C) Time course of GF-axis angles during the MD simulations of CytoD-bound actin. The initial structure was a crystal structure of F1^N13A^A_CytoD, with F1^N13A^ removed. The colored scatter points represent different runs. (D) Distribution of OD orientations from 300 – 400 ns in three runs of MD simulation of actin with F1^1-160^ (PDB code: 7W50). The complex also binds ADP and P_i_. The black points represent the initial orientation (14.9, −1.2). (E) Distribution of OD orientations from 300 – 400 ns in three runs of MD simulation of CytoD-bound actin. The black points represent the initial orientation (14.4, −1.4). (F) Comparison of actin conformations of F1^1-160^-bound actin and Cyto-D-bound actin, which were included in the peak bins of the heatmaps (D) and (E). The peak bins correspond to (10,-3) and (10,-8), respectively. The two conformations were pair-fitted using the ID domain (residue numbers: 147-166, 171-194, 205-229, 237-238, 251-322, and 327-337). Cyto-D binding (cyan) slightly opens the nucleotide-binding cleft relative to F1^1-160^ binding (green).

## Discussion

Our TIRF microscopic observations provide direct evidence that CytoD is an end capper that prevents both actin monomer association and dissociation at barbed ends. We look closer at the structural basis behind the inhibitory mechanism of CytoD on actin. Elongation inhibition by CytoD can be explained by simple steric hindrance. As proposed from CytoD-bound G-actin structures (11), CytoD binds to the hydrophobic cleft on the penultimate subunit, blocking monomer association by obstructing D-loop interaction. On the other hand, depolymerization inhibition by end cappers typically involves bridging the terminal subunit with adjacent actin subunits. For example, gelsolin links subunits within the same strand (31, 32), while CP connects subunits across strands (33, 34). Then how does CytoD, which can only interact with a single subunit, inhibit depolymerization? Recent cryo-EM studies established that the conformation of the two barbed end subunits is the F-form (35, 36). Thus, depolymerization at the barbed end involves a conformational transition from the F-form to the G-form of dissociating monomers, which disrupts D-loop mediated intra-strand interaction. Both the previous (11) and current crystal structures **(Fig. S2C-S2E)** demonstrate that CytoD shifts actin into a flatter conformation. This F-form-stabilizing effect of CytoD was further supported by our MD simulations **(Fig. 6C)**. We thus propose that CytoD inhibits actin depolymerization by binding to the hydrophobic cleft of barbed end subunits, indirectly stabilizing D-loop-mediated intra-strand interaction.

Previous reports showed that CytoD promotes the dimerization of Mg^2+^-ATP-actin at a 1:2 molar stoichiometry (12, 13). Based on the properties of CytoD discussed above, we propose that CytoD traps two actin molecules in the same orientation as adjacent subunits within a single strand of an actin filament (*i.e.,* a long-pitch dimer) through the following mechanism **(Fig. S4A)**. CytoD binds to the hydrophobic cleft of an actin monomer, inducing its flattening, which facilitates interaction with another actin monomer. The “upper subunit” cannot bind CytoD because its hydrophobic cleft is occupied by the D-loop of the “lower subunit”, leaving the upper subunit in the G-form, incompatible with further elongation. Without opposite strand subunits, the CytoD-induced long-pitch dimer is likely short-lived and further destabilized as ATP hydrolysis and subsequent P_i_ release convert the F-form lower subunit to ADP state, which cannot support dimer stability (13).

An intriguing phenomenon revealed by our kinetic analysis is the sustained elongation at rates slower than those of free actin, observed in polymerization assays with low concentrations of CytoD. Under these conditions, CytoD functions as a “leaky capper”, repeatedly associating with and dissociating from the barbed end, thereby permitting monomer association (**Fig. 7A left**, depicted as “Leakey cap). Conversely, in depolymerization assays and polymerization assays performed in the presence of high concentrations of CytoD, it acts as a stable barbed end capper over extended periods. What explains the conflicting observations regarding the fast versus slow dissociation of CytoD from the barbed end? A possible explanation for this may be due to the presence of two distinct capping states. A superposition of our CytoD-bound F-form actin structure onto the filament barbed end structure indicates that CytoD can bind without any steric clashes, even to the penultimate actin subunit **(Fig. S4B)**. This suggests two possible capping states for CytoD: a fully capped state, where both strands of the barbed end are capped by CytoD **(Fig. 7A**, bottom left corner), and a single-strand capped state, where only one strand is capped **(Fig. 7A**, middle left; the specific capped strand in not distinguished). CytoD dissociation from a fully capped barbed end likely occurs in two sequential steps: 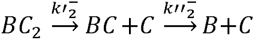 (*B* and *C* represent the barbed end and CytoD, respectively, while 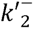 and 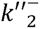 represent the dissociation rates for the first and second steps, respectively). We reanalyzed depolymerization assay data **(Fig. 1F)**, collected at 25 nM CytoD where both strands were capped, by applying this sequential dissociation model **(SI Appendix)**. The analysis identified two dissociation rates: 0.010 s⁻¹ (t_1/2_ = 70 s) and 0.080 s⁻¹ (t_1/2_ = 8.7 s, **Fig. S1F**). Although our analysis could not definitively assign these rates to specific events (k^-^_2_’ or k^-^_2_’’, **Fig. 7A** bottom left corner and middle left, respectively), it is reasonable to hypothesize that the faster dissociation rate corresponds to the second event, in which CytoD dissociates from the single-strand capped state (*i.e.*, k^-^_2_’’ = 0.080 s⁻¹). This interpretation aligns with the rapid dissociation observed at low CytoD concentrations, where single-strand capping is expected to predominate due to limited CytoD availability. Conversely, the slower dissociation rate likely corresponds to the first event, reflecting dissociation from the fully capped state (k^-^_2_’ = 0.010 s⁻¹). Our kinetic analysis thus supports a model in which CytoD stably caps the barbed end when both strands are simultaneously bound but provides only transient stabilization when binding occurs on a single strand. This dual binding mechanism reconciles the observed differences in dissociation kinetics and highlights the dynamic nature of CytoD capping at the barbed end.

**Figure 7.**
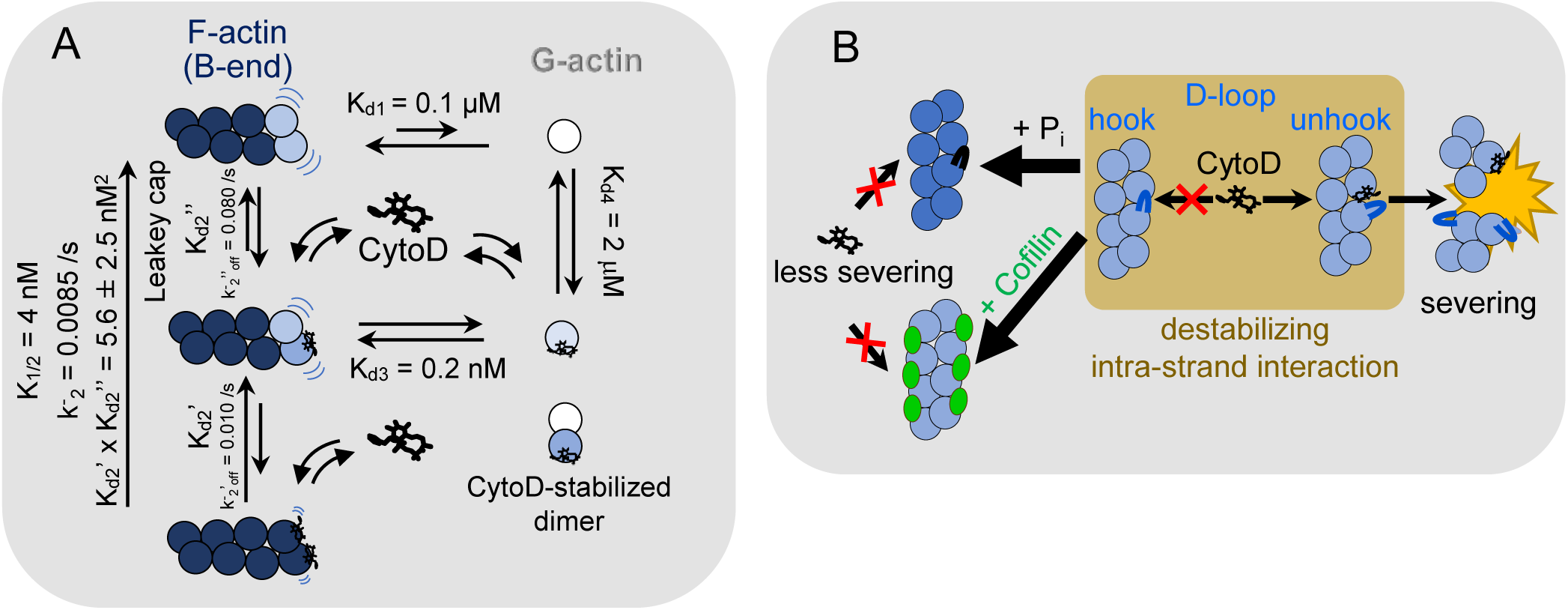
Mechanism of CytoD-induced inhibition of actin dynamics through capping and severing. (A) Barbed end capping by CytoD. The rates and equilibrium constants measured in this study, combined with previous data, provide a comprehensive view of the CytoD-dependent inhibition cycle of actin polymerization. A prior study reported the critical concentrations of actin bound to ATP (K_d1_) (26). CytoD binds to the barbed end in two sequential steps: initial binding to a free barbed end (K_d2_’’), followed by the binding of a second CytoD to a barbed end already bound by one CytoD (K_d2_’). Our elongation assays revealed that the moderate association and dissociation rates of CytoD at the barbed end leading partial capping, referred to as a “leaky cap”, where K_d2_’’ > K_d2_’. These dissociation constants must satisfy the relationship K_d2_’’ × K_d2_’ = 5.6 nM^2^, as determined in Fig 1D (SI Appendix). The difference between K_d2_’’ and K_d2_’ arises from the dissociation rates of CytoD from the barbed end: slow (k^-^_2_’= 0.010 s^-1^) and fast (k^-^_2_’’ = 0.080 s^-1^) (Fig. S1F and SI Appendix). CytoD also binds free actin monomers (G-actin) with relatively low affinity (K_d4_ = 2 μM) (9). The CytoD:G-actin complex can assemble with another G-actin, forming a longitudinal dimer in which the hydrophobic cleft of the upper subunit is occupied by the D-loop of the lower subunit. Kinetic analysis completed the thermodynamic square, showing that the binding affinity of CytoD-bound G-actin (K_d3_ ≈ 0.2 nM) is 1000 times tighter than actin alone (K_d1_ = ∼0.1 μM). CytoD-bound actin adopts the F-form, favorable for polymerization. However, this reaction remains minor due to the low affinity of CytoD for G-actin (K_d4_). (B) CytoD-induced severing of actin filaments. CytoD binding also promotes filament severing by destabilizing intra-strand interactions. Fluctuations in the D-loop expose the CytoD binding site without disrupting the overall filament structure. When CytoD binds to an actin subunit in the filament, the D-loop of the subunit below disengages from the upper subunit (“unhooking”), leading to severing. Both inorganic phosphate (P_i_) and cofilin inhibit CytoD-induced severing: P_i_ stabilizes intra-strand interactions and reduces subunit fluctuations, while cofilin facilitates a conformational change of actin to C-form, which is likely unfavorable for CytoD binding.

The proposed mechanism aligns with our structural data, demonstrating that the binding affinity for CytoD is highly sensitive to actin conformation. Structural flexibility at the filament ends likely allows actin to deviate from the optimal ligand-binding conformation, the F-form. Intrinsic structural fluctuations at the barbed end are likely unfavorable for CytoD binding, as CytoD has a low affinity for G-form actin (**Fig. 7A** right, K_d4_= 2 μM) (9). Additionally, subtle local conformational changes, such as a downshift of the Pro-rich loop, may further weaken the affinity for CytoD. Stable binding of CytoD to one strand is likely hindered by the wobbling of the adjacent free strand (**Fig. 7A** middle left). In this way, the structural fluctuations inherent to the barbed end confer cooperativity to CytoD binding across both strands, which would otherwise appear independent. Collectively, our results demonstrate that CytoD inhibits actin polymerization dynamics not only by sterically blocking the binding site but also by harnessing the intrinsic structural flexibility and fluctuations of actin.

The stoichiometry of CytoD binding to the actin filament has been a long-standing subject of debate (2). Although we did not directly observe this, our structural and kinetic results strongly support the conclusion that two CytoD molecules can bind simultaneously to a barbed end. This awaits confirmation through future cryo-EM observations of CytoD-capped filaments.

Our observation confirmed directly that CytoD can sever actin filaments. While this may not be surprising, as earlier studies have reported the filament-severing activity of other small compounds targeting actin’s hydrophobic cleft (37–39). These compounds, including CytoD, likely share a common mechanism when severing filaments, destabilizing intra-strand interaction through a competition with the D-loop (**Fig. 7B** surrounded by square). The significance of our findings lies in directly demonstrating the severing activity of CytoD, one of the most widely used toxins. We found that adding P_i_ suppresses CytoD-mediated severing of ADP-actin filament (**Fig. 7B** upper left). P_i_ acts as a universal inhibitor of severing molecules, including cofilin (40, 41) and gelsolin (42). CytoD (5) and gelsolin (43) target the actin’s hydrophobic cleft, which is occupied by the D-loop in the filament. P_i_ likely prevents severing by stabilizing the actin conformation in the F-form and reducing the frequency with which the D-loop dissociates. We also found that filaments decorated with cofilin are resistant to severing by CytoD (**Fig. 7B** bottom left). This observation is likely due to cofilin’s binding to actin filaments, which prompts a transition to a unique conformation (C-form) (44) that appears unfavorable for CytoD binding. This, in turn, may explain why rapid filament severing driven by a three-component system involving cofilin could be inhibited by CytoD (20). Severing through competition with the D-loop is unlikely to occur in the cofilin-bound filaments in which the D-loop is dissociated (44).

Given the severing that occurs with several µM of CytoD, a range typically used in cellular experiments, filament cleavage should be considered when interpreting the effects of CytoD on the cellular actin cytoskeleton. Indeed, observations have been reported that can be explained by an increase in short filaments resulting from CytoD-induced cleavage (45, 46). Importantly, CytoD-induced severing does not occur uniformly; unstable filaments, such as aged ADP-actin filaments, will be targeted selectively, while filaments decorated with cofilin or other actin-binding proteins may be protected.

## Materials and Methods

### Protein purification and labeling

Actin was prepared from acetone powder from rabbit (TIRF experiments) or chicken (crystallization) skeletal muscle by one cycle of polymerization and depolymerization (47). We labeled actin on lysine with AF488 NHS ester (Lumiprobe, #21820, USA) as described (18). The expression vector for the fragmin (*Physarum polycephalum*) F1 domain carrying an Asn13Ala mutation (F1^N13A^) was generated using the KOD plus mutagenesis kit (Toyobo), with the wild-type F1 expression vector (F1; GST-tag at the N-terminus (27)) serving as the template. Both F1 and F1^N13A^ were expressed in *Escherichia coli* BL21(DE3) and purified as previously described (48). Cytochalasin D (Sigma-Aldrich) was dissolved in Dimethyl sulfoxide at a concentration of 2 mM and stored at −20°C, and diluted into polymerization buffer. Human cofilin-1 was expressed in *E. coli* BL21 (DE3) or C43 (DE3) for 24 h at 15 °C with induction conditions of OD_600_ = 0.4 and IPTG = 1 mM. Cell pellets were resuspended in a buffer consisting of 50 mM Tris-HCl pH 8.0, 300 mM NaCl, 50 mM imidazole-HCl pH 8.0 and lysed by sonication. The lysate was clarified by centrifugation and the protein was purified using a HisTrap HP column (Cytiva), followed by a HiLoad 26/600 Superdex 200 pg. column (Cytiva) equilibrated with a buffer consisting of 20 mM HEPES (pH 7.5), 150 mM NaCl, 0.5 mM EGTA at 4 °C.

### TIRF assay to observe individual actin filaments

A previous paper (18) describes the assay with the method to prepare the observation chamber. Briefly, coverslips were cleaned with KOH and double distilled water wash using sonicator and air-dried. The chamber was formed by placing one slide glass and the cleaned coverslip that were attached by double sided tape. The chamber was treated with 50 nM N-ethylmaleimide (NEM) myosin, which is inactivated myosin prepared from skeletal muscle (49) for 90 seconds at room temperature. BSA was then flowed into the chamber and incubated for 90 seconds to block nonspecific binding of actin on the glass surface. NEM-myosin and BSA were diluted in a buffer containing 600 mM KCl and 50 mM Tris-HCl, pH 8.0. Actin (final concentration was 0.7 μM, unless otherwise noted) was then mixed with TIRF buffer (100 mM KCl, 2 mM MgCl_2_, 0.5 mM EGTA, 10 mM Tris-HCl, pH 8.0, 25 mM DTT, 2 mM ATP, 0.5 % (wt/vol) methylcellulose [1500 cP] (18) with 10 mM sodium ascorbate as an oxygen scavenger) in the sample tube to initiate the polymerization, then immediately loaded into the chamber. Observations were carried out at 20-22 °C. To demonstrate CytoD effect on actin association, actin with a range of concentrations of CytoD in TIRF buffer was loaded after confirming which end is fast growing end of actin filaments on the slide. To observe depolymerization inhibition and severing induced by CytoD, TIRF buffer containing only various concentrations of CytoD was loaded and free actin monomer in the chamber was washed. Samples were kept in the dark except when images were taken at every 5 seconds. To prepare fully decorated actin with cofilin (cofilactin), 3 μM cofilin in TIRF buffer with imidazole-HCl, pH 6.6 instead of Tris-HCl, pH 8.0 was introduced into the observation chamber, which already contained polymerized actin (final concentration, 0.4 μM). The mixture was then incubated for approximately 5 minutes at 20–22 °C. No NEM-myosin was added to this chamber, so actin filaments were anchored to the glass surface through random adhesion.

### Image acquisition and analysis

AF488-labeled actin filaments were observed using a Nikon Ti2-E inverted microscope equipped with a 100× objective lens and a 488 nm laser. Images were captured with 250 or 500 ms exposures on an iXON EMCCD camera (Andor iXon, UK), with a pixel size of 0.16 μm in the sample plane. Actin filament lengths were measured using ImageJ (NIH) with plugins generously provided by Jeff Kuhn (19).

### Crystallization and structural determination

Our initial attempt to obtain the CytoD-bound F1A structure by soaking apo crystals with CytoD was unsuccessful. Although the soaked crystals diffracted well, no density assignable to CytoD was observed. Therefore, F1^N13A^ (fragmin residues 1-160, with Asn13 replaced by Ala) was employed for crystallization, as this mutation in FNEX was expected to reduce interference with CytoD binding to actin. One microliter of protein solution containing F1^N13A^ and actin (165 µM each), supplemented with 2.5 mM CytoD, was added to an equal volume of reservoir solution (17% w/v PEG3350 and 20 mM Na_2_HPO_4_) on a cover slip and streaked with a needle dipped into a seed solution containing crushed apo crystals. The coverslips with seeded drops were placed onto a 24-well hanging drop plate (Hampton Research) and incubated at 20°C against the reservoir solution (500 µl). After one week of incubation with repeated streak seeding, a tiny rod-shaped crystal (diameter of ∼0.02 mm) appeared in the drop. The crystal was cryo-protected by briefly soaking it in the reservoir solution supplemented with 15% (v/v) ethylene glycol and then flash-cooled in a cold nitrogen stream. X-ray diffraction measurements were performed on beamline BL2S1 at the Aichi Synchrotron Radiation Center (50) with a wavelength of 1.12 Å at −183°C. The collected dataset was processed with XDS (51). The initial phase was obtained by molecular replacement with Molrep (52) using the apo structure (PDB code 7W50) as the search model. Restrained refinement was performed with Refmac5 (53) and Phenix_refine (54), and model inspection was carried out using Coot (55). After deleting FNEX residues that lacked associated electron density, a large blob of unassigned electron density was visible between subdomains 1 and 3 of actin, which was modeled as CytoD. The position and orientation of CytoD were almost identical to those observed in CytoD-bound G-form actin structures (3EKS and 3EKU;(11)). All structural figures were prepared using PyMOL (https://www.pymol.org/). Data collection and refinement statistics are summarized in **Table S1**.

### MD simulation

MD simulations were performed using the GROMACS 2023.1 (56) with CHARMM36-jul2021 force field (57) and TIP3p force field (58) on FUJITSU PRIMERGY CX2570 M5 computer at Nagoya University Information and Communications. Force field parameters for CytoD and Pi were generated using SwissParam (59). F1^N13A^A_CytoD (in this study) and F1^160^-A (PDB code: 7W51) were used as initial structures. The missing regions, the N-terminus and the D-loop of the initial crystal structures were reconstructed based on other actin crystal structures (29). The initial proteins were solvated in a rectangular box with a minimum distance of 1.5 nm from the protein to the box edge, using both crystal water molecules and additional water molecules generated by GROMACS. Potassium and chloride ions (100 mM) were added for the net charge of the system. Electrostatic interactions were calculated using the particle-mesh Ewald algorithm (60), and all bond lengths were constrained using the linear constraint solver algorithm (61). The system was maintained at 300 K and 1 atm using a v-rescale thermostat (62) and a Parrinello–Rahman barostat (63), respectively. Following energy minimization of with the steep distant method, the system was equilibrated under constant volume and temperature for 1 ns with position restraints for the heavy atoms of the protein, Mg^2+^, and ligand, followed by equilibration for 1 ns under constant pressure and temperature. Subsequently, 400 ns MD productions were conducted. The resulting traces were analyzed using GROMACS, Scilab (https://www.scilab.org/), and PyMOL.

## Supporting information

revised SI figures

## Acknowledgments

X-ray diffraction measurements were performed on beamline BL2S1 at the Aichi Synchrotron Radiation Center. We thank the beamline staff for their technical assistance. This work was supported by Japan Society for the Promotion of Science grants KAKENHI 16K17708, 20K06522, and 23K05666 (to S.T.), 17K07373 (to S.T., T.O., and I.F.), 22K06172 (to S.T. and T.O.), 24K08218 (to I.F.), Iketani Science and Technology Foundation (0361200-A) (to I.F.).

