## Supplementary material for "Microscopic and structural observations of actin filament capping and severing by Cytochalasin D": revised SI figures

#### Supporting Information for
### Microscopic and structural observations of actin filament capping and severing by Cytochalasin D

Takahiro Mitani, Shuichi Takeda*, Toshiro Oda, Akihiro Narita, Yuichiro Maéda[4], Hajime Honda and Ikuko Fujiwara†

**This PDF file includes:**

Supporting text
Figures S1–S4
Tables S1 and S2
Legends for Movies S1–S3
SI References

**Other supporting materials for this manuscript include the following:**

Movies S1 and S2

**Supporting Information Text**

**Method 1: Estimation of the combined equilibrium dissociation rates $K'_{d2} \times K''_{d2}$**

Our used model is below.

$B \xrightarrow{K''_{d2}} BC_1 + C \xrightarrow{K'_{d2}} BC_2$

where $B$ and $C$ represent actin at the barbed end and CytoD, respectively. $BC_2$ and $BC$ represent barbed ends bound two CytoD molecules and one CytoD molecule, respectively.

$$K''_{d2} = \frac{[\text{Free B}][\text{Free C}]}{[BC_1]}$$

$$K'_{d2} = \frac{[BC_1][\text{Free C}]}{[BC_2]}$$

$$[BC_2] = \frac{[\text{Free B}][\text{Free C}]^2}{K'_{d2} \times K''_{d2}}$$

where. $[Total\ B] = [Free\ B] + [BC_1] + [AB\ ]$ Then, the following equation is applied to Fig. 1D to estimate $K'_{d2} \times K''_{d2}$.

$$[BC_2] = \frac{[\text{Free C}]^2}{K'_{d2} \times K''_{d2} + [\text{Free C}]^2}$$

**Method 2: sequential dissociation reaction kinetics.**

Our used model is below (SI ref1).

$BC_2 \xrightarrow{k'^-_2} BC + C \xrightarrow{k''^-_2} B + 2C$

where $B$ and $C$ represent actin at the barbed end and CytoD, respectively. $BC_2$ and $BC$ represent barbed ends bound two CytoD molecules and one CytoD molecule, respectively. The dissociation rates for first and second reactions are $k'^-_2$ and $k''^-_2$, respectively. Differential equations are as follows.

$$\frac{d[BC_2]}{dt} = -k'^-_2[BC_2]$$

$$\frac{d[BC]}{dt} = k'^-_2[BC_2] - k''^-_2[BC]$$

$$\frac{d[B]}{dt} = k''^-_2[BC]$$

The fraction of $[BC_2]$ and $[BC]$ decrease over time

$$[BC_2] = [BC_2]_0 e^{-k'^-_2 t}$$

Here, $[BC_2]_0 = [BC_2] + [BC] + [B\ ]$

$$[BC] = \frac{k_2^{\prime -}[BC_2]_0}{k_2^{\prime\prime -} - k_2^{\prime -}}\left(e^{-k_2^{\prime -}t} - e^{-k_2^{\prime\prime -}t}\right)$$

$$[B] = [BC_2]_0\left\{1 + \frac{1}{k_2^{\prime -} - k_2^{\prime\prime -}}\left(k_2^{\prime\prime -}e^{-k_2^{\prime -}t} - k_2^{\prime -}e^{-k_2^{\prime\prime -}t}\right)\right\}$$

Thus, the equation to fit Fig. 1F is below.

$$[BC_2]_0 - [B] = \frac{[BC_2]_0}{k_2^{\prime -} - k_2^{\prime\prime -}}\left(k_2^{\prime -}e^{-k_2^{\prime\prime -}t} - k_2^{\prime\prime -}e^{-k_2^{\prime -}t}\right)$$

$[BC_2]_0 = 1$ was applied for actual data fitting, since the data is normalized.

For more detail,

$$\frac{d[B]}{dt} = k_2^{\prime\prime -}[BC]$$

$$\frac{d[B]}{dt} = k_2^{\prime\prime -}[BC] = k_2^{\prime\prime -}\frac{k_2^{\prime -}[BC_2]_0}{k_2^{\prime\prime -} - k_2^{\prime -}}\left(e^{-k_2^{\prime -}t} - e^{-k_2^{\prime\prime -}t}\right)$$

$$\frac{k_2^{\prime\prime -} - k_2^{\prime -}}{k_2^{\prime -}k_2^{\prime\prime -}}d[B] = \left(e^{-k_2^{\prime -}t} - e^{-k_2^{\prime\prime -}t}\right)dt$$

$[A] = 0$ when $t = 0$, thus upon integration,

$$\frac{k_2^{\prime\prime -} - k_2^{\prime -}}{k_2^{\prime -}k_2^{\prime\prime -}}\frac{[B]}{[BC_2]_0} = \int_0^t\left(e^{-k_2^{\prime -}t} - e^{-k_2^{\prime\prime -}t}\right)dt = \left[-\frac{1}{k_2^{\prime -}}e^{-k_2^{\prime -}t} + \frac{1}{k_2^{\prime\prime -}}e^{-k_2^{\prime\prime -}t}\right]$$

$$= -\frac{1}{k_2^{\prime -}}e^{-k_2^{\prime -}t} + \frac{1}{k_2^{\prime\prime -}}e^{-k_2^{\prime\prime -}t} + \frac{1}{k_2^{\prime -}} - \frac{1}{k_2^{\prime\prime -}}$$

Thus,

$$\frac{k_2^{\prime\prime -} - k_2^{\prime -}}{k_2^{\prime -}k_2^{\prime\prime -}}\frac{[B]}{[BC_2]_0} = \frac{1}{k_2^{\prime -}k_2^{\prime\prime -}}\left(-k_2^{\prime\prime -}e^{-k_2^{\prime -}t} + k_2^{\prime -}e^{-k_2 t} + k_2^{\prime\prime -} - k_2^{\prime -}\right)$$

$$\frac{k_2^{\prime\prime -} - k_2^{\prime -}}{k_2^{\prime -}k_2^{\prime\prime -}}\frac{[B]}{[BC_2]_0} = \frac{1}{k_2^{\prime -}k_2^{\prime\prime -}}\left(-k_2^{\prime\prime -}e^{-k_1 t} + k_2^{\prime -}e^{-k_2^{\prime\prime -}t} + k_2^{\prime\prime -} - k_2^{\prime -}\right)$$

$$[B] = [BC_2]_0\left(1 + \frac{1}{k_2^{\prime -} - k_2^{\prime\prime -}}\left(k_2^{\prime\prime -}e^{-k_2^{\prime -}t} - k_2^{\prime -}e^{-k_2^{\prime\prime -}t}\right)\right)$$

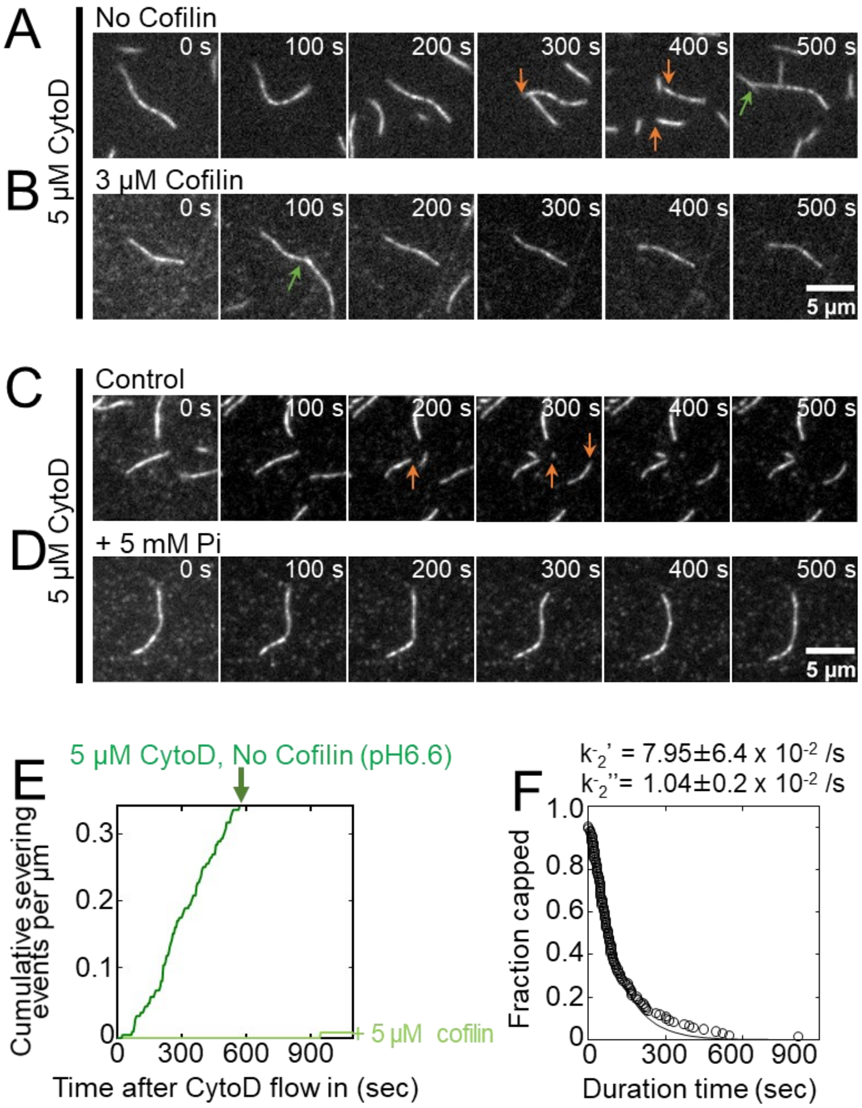

**Fig. S1.** Severing inhibition by ADF/cofilin or $P_i$.

(A and B) Still images of actin filaments after washing out free actin monomer with the buffer (t= 0) containing 5 μM CytoD in the presence of 0 μM (A) or 3 μM cofilin (B). Arrows show severed points (orange) and filaments overlapping (green). Time indicates when free actin monomer was washed out with the buffer containing 5 μM CytoD (t= 0). Scale bars, 5 μm. (C and D) Same assay as (A and B), washing out free actin monomer with 5 μM CytoD (t= 0) as control (C) or only the buffer contains 5 mM $P_i$ instead of cofilin (D). Scale bars, 5 μm. (E) Cumulative severing events on actin filaments per μm over time in the absence (dark green) or presence (green) of 5 μM CytoD at the polymerization buffer (pH 6.6). CytoD severed actin filaments even under low pH, which was suppressed in the presence of 5 μM cofilin (green, same as Fig. 3H). Actin filaments were not anchored with NEM-myosin, because cofilactin does not bind it. The increased severing of cofilactin may be due to the fluctuation of the filament. (F) Duration time of capping for each actin filament plotted as τ. The $k_{-2}$' and $k_{-2}$'' values obtained from double exponential fit were $7.95 \pm 6.4 \times 10^{-2}$ and $1.04 \pm 0.20 \times 10^{-2}$ $s^{-1}$, estimated from 116 filaments in two independent experiments.

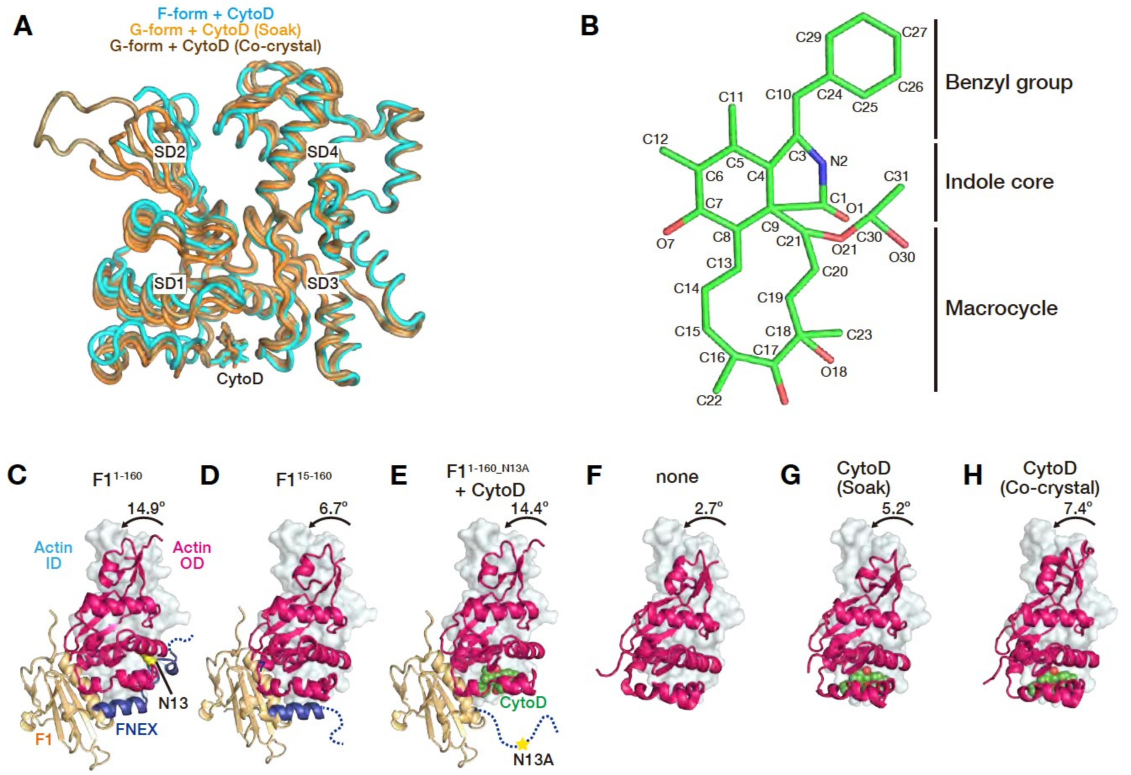

**Fig. S2.** CytoD binding to actin and its effect on actin conformation

(A) Comparison of the Cyto-D bound F-form (cyan; this study) and G-form actin (orange; soaked, PDB 3EKU and brown; co-crystallized, 3EKS) (2) reveals that CytoD occupies the same location between SD1 and SD3 of actin. (B) The chemical structure of CytoD with numbered atoms (hydrogens omitted). (C-H) Actin conformation. Various actin structures complexed with fragmin F1 and/or CytoD are shown from the OD side. The twisting angle of the OD (pink) relative to the ID (white) around the GF axis (see Fig. 6A) is quantified. The G-actin structure (PDB 1J6Z) (3) is used as the reference with a twisting angle set as 0°. (C) F1$^{1\text{-}160}$-bound actin (7W50) (4). Actin adopts an F-form conformation, stabilized by FNEX interactions at both the bottom and back. (D) F1$^{15\text{-}160}$-bound actin (chains A and B, PDB 7W52) (4). The absence of Asn13 disrupts back-side actin binding, resulting in a G-form conformation. (E) F1$^{1\text{-}160_N13A}$ and CytoD-bound actin (this study). Despite the lack of FNEX interaction, actin adopts an F-form conformation. This indicates that CytoD (spheres) stabilizes the flat actin conformation, aided by a crystal lattice designed for aligning F1-bound F-form actin. Crystals did not form from F1$^{1\text{-}160_N13A}$ and actin in the absence of CytoD. (F) CytoD-free actin (PDB 2HF4) (5). Recombinantly expressed, non-polymerizable actin adopts a typical G-form conformation in the crystal. (G) CytoD-soaked actin (PDB 3EKU) (2). Actin exhibits slight flattening (2.5°) compared to the CytoD-free structure. (H) CytoD co-crystallized actin (PDB 3EKS) (2). The actin structure is further flattened (2.2°) compared to the soaked structure, supporting previous suggestions that CytoD stabilizes a flatter actin conformation actin (2).

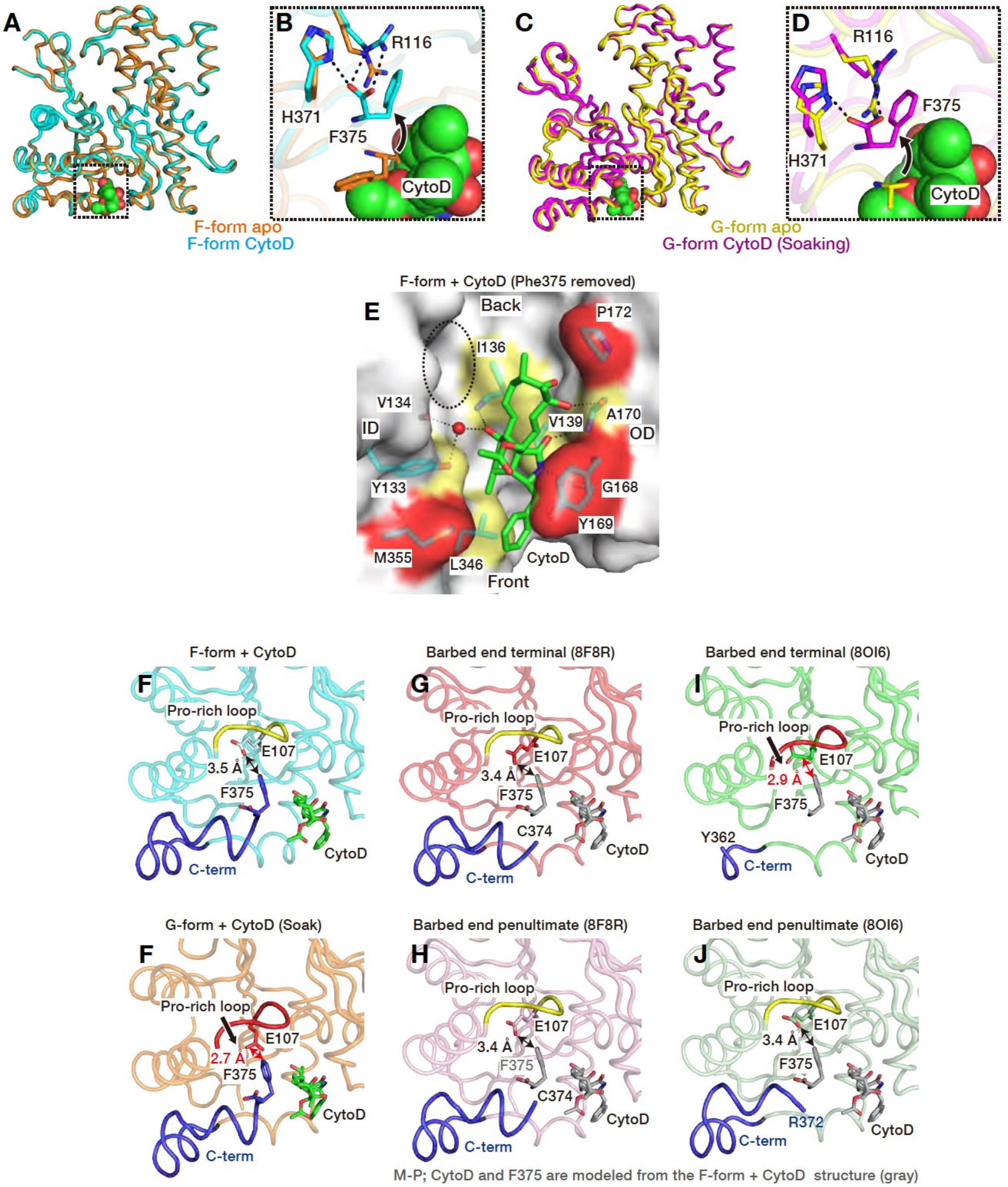

Figure legend in the next page

**Fig. S3.** Phe375 in CytoD-bound actin structures.
(A-D) CytoD-induced Phe375 repositioning in the F-form (A-B) and G-form (C-D) actins. The coordinates are as follows: F-form without CytoD (PDB 7W50), F-form with CytoD (this study), G-form without CytoD (2HF4), G-form with CytoD (soaking; 3EKU). Curved arrows indicate the flipping of Phe375 to avoid CytoD. In the flipped Phe375 conformation, two carboxyl oxygens of Phe375 electrostatically interact with the side chain nitrogen of Arg116 and His371. A cation-π interaction is formed between the phenyl ring of Phe375 and the guanidinium group of Arg116.
(E) Phe375-removed actin model. CytoD-bound F-form actin (surface representation) from which Phe375 is removed. This deletion model indicates that Phe375 (whose occupation in the intact structure is shown by a dotted circle) is indispensable in forming a hydrophobic pocket accommodating CytoD.
(F-J) Models of CytoD interaction with the filament barbed end actin subunits. As a representative of potential clashes between CytoD and actin residues, the distance between Phe375 and Glu107 is shown in the following structures: (F) CytoD-bound F-form actin, (F) CytoD-bound G-form actin (soaked), (G-J) Cryo-EM structure of the free filament barbed end (8F8R) (6); terminal (G) and penultimate (H) actin subunits, (I-J) Cryo-EM structure of the phalloidin-stabilized filament barbed end (8OI6) (7); terminal (I) and penultimate (J) actin subunits. Actin residues from

Thr358 to the C-terminus (indicated) are colored blue. The Pro-rich loop (residues 108-112) adopting a conformation observed in the filament interior subunits is colored yellow. In panels G-J, Phe375, and CytoD, derived from the CytoD-bound F-form structure, are modeled in gray sticks. In G-form actin (F), and in the terminal subunit of one barbed end structure (I), the Pro-rich loop is downward-shifted (red). The actin C-terminus is not folded in barbed end structures, suggesting that Phe375 does not obstruct CytoD incorporation. However, as shown above, for tight binding with CytoD, Phe375 must participate in the hydrophobic pocket to stabilize the macrocycle ring, which seems unaffected by its surrounding residues. An exception is the terminal actin subunit in a barbed end model (8OI6), where the Pro-rich loop is downward-shifted, resembling its position in G-actin (Fig. S3I). This suggests that CytoD needs to push up the Pro-rich loop to bind to that subunit, as incoming monomers acting on the penultimate subunit (7).

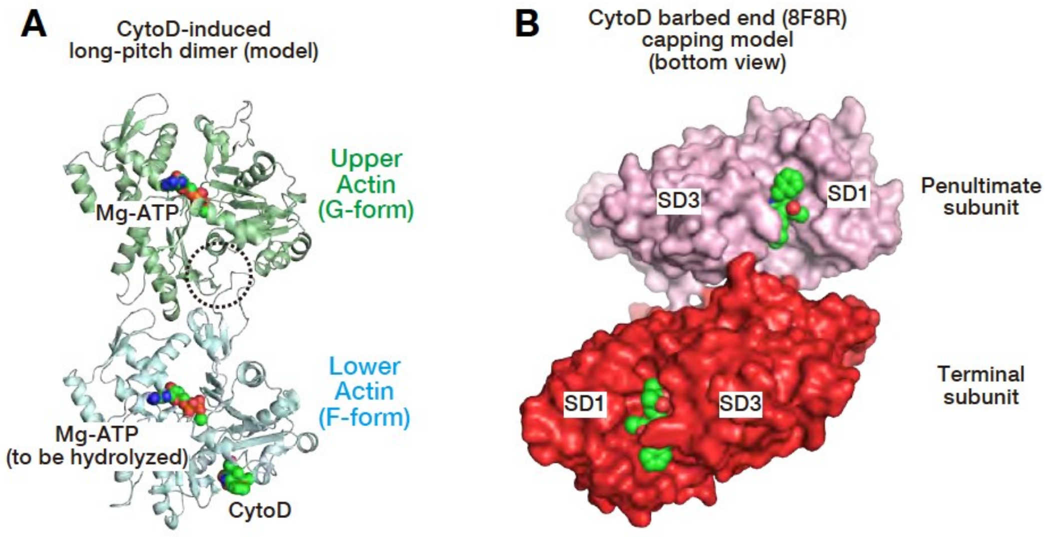

**Fig. S4.** CytoD-bound actin models.
(A) A model of CytoD-induced long-pitch actin dimer. See discussion for details. (B) A model of CytoD capping the barbed end. Our CytoD-bound F-form actin structure was superimposed onto the two barbed end subunits in a cryo-EM structure (8F8R), viewed from the bottom of the penultimate subunit (pink) to highlight the inter-subunit interface with the terminal subunit (red). Actin subdomains are labeled for reference. In this model, CytoD readily binds to the terminal actin subunit. Moreover, CytoD binding to the penultimate subunit appears unaffected by the terminal actin subunit. This model yields two key insights: (2) two CytoD molecules can bind to a single actin filament, and (3) CytoD neither stabilizes nor destabilizes inter-strand interactions between barbed end actin subunits. However, we cannot exclude the possibility that CytoD binding, particularly to the penultimate subunit, induces local structural rearrangements within that subunit, thereby enhancing its interactions with the terminal subunit.

Table S1. **Data collection and refinement statistics**

| | $F1^{N13A}A$_CytoD (PDB 9L2N) |
|---|---|
| **Data collection** | |
| Space group | $P2_12_12_1$ |
| Cell dimensions | |
| *a*, *b*, *c* (Å) | 57.2, 90.8, 114.6 |
| *α, β, γ* (°) | 90, 90, 90 |
| Resolution (Å) | 48.44-1.70 (1.80-1.70) |
| No. unique reflections | 65980 (10466) |
| $R_{\text{merge}}$ | 0.112 (0.854) |
| $R_{\text{meas}}$ | 0.143 (1.09) |
| $R_{\text{pim}}$ | 0.0875 (0.670) |
| $CC_{1/2}$ | 0.99 (0.63) |
| $I$ / σ$I$ | 8.75 (1.49) |
| Completeness (%) | 99.5 (98.3) |
| Redundancy | 4.7 (4.7) |
| **Refinement** | |
| $R_{\text{work}}$ / $R_{\text{free}}$ | 0.175/0.204 (0.269/0.281) |
| No. atoms | |
| Protein | 3981 |
| Ligand/ion | 90 |
| Water | 870 |
| *B*-factors | |
| Protein | 19.4 |
| Ligand/ion | 22.1 |
| Water | 33.4 |
| R.m.s. deviations | |
| Bond lengths (Å) | 0.011 |
| Bond angles (°) | 0.96 |
| Ramachandran | |
| Favored (%) | 97.9 |
| Outliers (%) | 0 |
| All-atom Clashscore | 2.11 |
| Molprobity score | 1.00 |

Table S2. **CytoD contacting residues**

| Residues | Soaked G-form (3EKU) | Co-crystallized G-form (3EKS) | F-form (This study) |
|---|---|---|---|
| Tyr133 | 17.80 | 21.66 | 26.05 |
| Ile136 | 24.35 | 25.50 | 23.88 |
| Val139 | 17.91 | 19.08 | 22.76 |
| Tyr143 | 20.93 | 21.44 | 20.54 |
| **Tyr169** | **64.52** | **68.56** | **69.28** |
| Ala170 | 20.35 | 21.97 | 26.71 |
| **Pro172** | **34.62** | **39.98** | **34.46** |
| Leu346 | 20.88 | 18.65 | (10.58) |
| **Met355** | **45.53** | **37.99** | **41.96** |
| Cys374 | (14.49) | 21.50 | 21.51 |
| **Phe375** | **45.74** | **35.77** | **35.43** |

All actin residues with a buried surface area (BSA; Å²) greater than 15 Å² upon CytoD binding are listed (values less than 15 Å² are enclosed in parentheses). Values exceeding 30 Å² are emphasized in bold. The BSA values were calculated using PDBePISA (8).

**Movie S1 (separate file).** CytoD inhibits actin depolymerization. The left and right movies show the depolymerization of filaments in the presence of 0 nM and 25 nM CytoD, respectively. The first 200 seconds are polymerization conditions followed by a sequence of solution changes. The colored arrows match those in the corresponding movie stills shown in Figure 1 and move in line with the growing barbed end. Images were collected at a rate of one frame every 5 s and played back in real time at 50× magnification.

**Movie S2 (separate file).** CytoD inhibits actin polymerization. The three movies show filament growth in the presence of 0 (left), 0.5 (middle), and 5 (right) nM of CytoD, respectively. The order of the solution exchanges is indicated. The colored arrows match the colored arrows in the corresponding the graph presented in Fig. 2C, and they move in register with the growing barbed end. Images were collected at a rate of 1 frame every 5 s and are played back at 50× real time.

**Movie S3 (separate file).** CytoD severs actin filaments. Movie shows filament severing in the presence of 10 µM CytoD. Actin was first polymerized and then CytoD was added. Arrows indicate the position and moment of severing.
